# Evaluation of Enhanced Learning Techniques for Segmenting Ischaemic Stroke Lesions in Brain Magnetic Resonance Perfusion Images using a Convolutional Neural Network Scheme

**DOI:** 10.1101/544858

**Authors:** Carlos Uziel Perez Malla, Maria del C. Valdes Hernandez, Muhammad Febrian Rachmadi, Taku Komura

**Affiliations:** University of Edinburgh, School of Informatics, Edinburgh, EH8 9AB, UK; University of Edinburgh, Department of Neuroimaging Sciences, Edinburgh, EH16 4SB, UK

**Keywords:** ischaemic stroke, medical image analysis, deep learning, computer vision, convolutional neural networks, deepmedic

## Abstract

Magnetic resonance (MR) perfusion imaging non-invasively measures cerebral perfusion, which describes the blood’s passage through the brain’s vascular network. Therefore it is widely used to assess cerebral ischaemia. Convolutional Neural Networks (CNN) constitute the state-of-the-art method in automatic pattern recognition and hence, in segmentation tasks. But none of the CNN architectures developed to date have achieved high accuracy when segmenting ischaemic stroke lesions, being the main reasons their heterogeneity in location, shape, size, image intensity and texture, especially in this imaging modality. We use a freely available CNN framework, developed for MR imaging lesion segmentation, as core algorithm to evaluate the impact of enhanced machine learning techniques, namely data augmentation, transfer learning and post-processing, in the segmentation of stroke lesions using the ISLES 2017 dataset, which contains expert annotated diffusion-weighted perfusion and diffusion brain MRI of 43 stroke patients. Of all the techniques evaluated, data augmentation with binary closing achieved the best results, improving the mean Dice score in 17% over the baseline model. Consistent with previous works, better performance was obtained in the presence of large lesions.

## 1 INTRODUCTION

Magnetic resonance imaging (MRI) has become a powerful clinical tool for diagnostics. Its application has been expanded to the evaluation of brain function through the assessment of a number of functional and metabolic parameters. One such parameter is cerebral perfusion, which describes the passage of blood through the brain’s vascular network. Amongst the several techniques used to measure cerebral perfusion (Fantini et al., 2016; Petrella and Provenzale, 2000), MRI is perhaps the most widely used due to its non-invasiveness. Thus, having great potential in becoming an important tool in the diagnosis and treatment of patients with cerebrovascular disease and other brain disorders. It measures cerebral perfusion via assessment of various hemodynamic measurements such as cerebral blood volume, cerebral blood flow, and mean transit time, from serial tissue tracer concentration measurements. These measurements are analysed in relation to their values in normal tissue regions (e.g. normal-appearing white matter). Therefore, the importance of estimating the location and extent of the abnormal region automatically.

Expert delineation is usually performed in the imaging modality that best displays the pathology while simultaneously evaluating other imaging modalities. The quality of this process depends on the expert’s experience, and suffers from intra- and inter-observer variability (Kamnitsas et al., 2017). Automated segmentation methods are not only necessary to provide the quantitative information needed to better support clinical decisions, but also to carry out large scale studies, with increased reliability and reproducibility, for which manual delineation is simply unattainable (Maier et al., 2017). Most of these algorithms use expert-labelled data to “learn” the pattern to be segmented until a certain level of accuracy is reached, and are expected to reproduce similar accuracy levels for new unlabelled data. Deep Learning algorithms, such as Convolutional Neural Networks (CNN), have risen in popularity due to their success on computer vision research (Krizhevsky et al., 2012). Though CNNs are typically used for multi-label image classification problems, they can also be employed for segmentation tasks by classifying each voxel according to the region they belong to (Kamnitsas et al., 2017).

In MR perfusion imaging, the pathologies’ appearance does not follow a clear pattern, which makes their detection far more difficult. Specifically ischaemic lesions can appear anywhere in the brain and their shape and signal intensities vary not only between disease stages but also within them (Maier et al., 2017). This variability increases with time from the stroke onset. Also, the intensity within the infarcted region is not necessarily homogeneous (Kamnitsas et al., 2017).

### 1.1 CNN Architectures for Brain Lesion Segmentation - DeepMedic

Specifically for the segmentation of brain lesions, different CNNs architectures have been evaluated(Guerrero et al., 2018; He et al., 2016). One of them(Guerrero et al., 2018) proposed a 2D CNN architecture for White Matter Hyperintensities (WMH) segmentation, and reported having achieved state of the art performance in differentiating them from ischaemic stroke lesions. However, by taking a 2D approach, it discards important spatial information, since did not take into account the volumetric nature of the data; and was only evaluated using structural MRI modalities, where lesions are homogeneous and easier to identify.

Using a 3D approach to manipulate Magnetic Resonance Imaging (MRI) data is not straightforward, as it requires significantly more computing power and memory than the 2D counterparts (Roth et al., 2014). The main factor that attempts against 3D segmentation is the slow inference process. This can be alleviated by taking advantage of dense inference (Sermanet et al., 2013), a property of full convolutional networks that avoids recomputing convolutions for overlapping image patches and thus reduces inference times. 3D CNN architectures have been used to segment pathologies, (Milletari et al., 2016; Brosch et al., 2016). However, DeepMedic (Kamnitsas et al., 2017) has emerged as the brain lesion segmentation CNN method for excellence, due to its availability, technical support and versatility, as it has been applied not only to segment hyperintense lesions (Rachmadi et al., 2018b), but also lesions with heterogeneous signal intensities (i.e. tumours) (Kamnitsas et al., 2017). It has a 3D CNN architecture of two pathways that uses dense-inference and adds a 3D fully connected Conditional Random Forest (CRF) as a final post-processing layer. By taking advantage of the dense inference, DeepMedic can be trained using image segments (i.e. image patches of size bigger than the network’s receptive field) to avoid recomputing convolutions of overlapping patches. Additionally, the dual pathway is used to compute both local and global (i.e. contextual) features at the same time by processing the same image at different scales. Finally, the CRF is used to remove false positives before returning the final results. DeepMedic reached the first position in the **I**schemic **S**troke lesion **S**egmentation (SISS) subchallenge of the **I**schemic **S**troke **LE**sion **S**egmentation (ISLES) 2015 challenge^1^.

Subsequent winners of the ISLES challenges have used other approaches. For example, whilst DeepMedic uses a traditional cross-entropy function (Kamnitsas et al., 2017), the winners of the ISLES 2017 challenge (Choi et al., 2017; Lucas and Heinrich, 2017), use a loss function based on Dice Similarity Coefficient (DSC) particularly designed for unbalanced data sets (Sudre et al., 2017). Also, (Choi et al., 2017) implement a spatial pyramid pooling layer (He et al., 2014), recently combined with an encoder-decoder (Chen et al., 2018b) to improve segmentation predictions. Spatial pyramid pooling guarantees a fixed output size for different sized inputs (He et al., 2014). This means that the network can process inputs at different scales, similarly to DeepMedic, while keeping the same output size. Dilated convolutions have also proven useful for enhancing the spatial resolution of the network and thus improving the performance for semantic segmentation (Chen et al., 2018a, 2017). These convolutional layers extend the field of view and thus can extract features at different scales.

### 1.2 Enhancing Learning Techniques

Variations in CNN architectures appear to show improvements in the segmentation of certain pathologies. However, these methods suffer a significant loss in performance when these changes are applied to datasets acquired with different imaging protocols, or using different sequences (i.e. task domain changes), they are applied to the assessment of different types of lesions caused by different pathology (e.g. the initial task being to segment tumour lesions, whilst the actual task is to segment ischaemic stroke lesions), or they are expected to perform tasks that are related to but not the same task they were trained for (e.g. lesion segmentation vs. lesion assessment).

There are several ways to enhance the performance of the CNN architectures without modifying the architecture itself. In general, they can be enumerated as follows: 1) pre-processing the input data, 2) modifying the input data by adding information derived from internal and external sources (i.e. data augmentation), 3) re-purposing a model trained for one task to perform a second related task (i.e. transfer learning), and 4) post-processing the output from the CNN.

#### 1.2.1 Pre-processing the Input Data

The importance of pre-processing the data has been highlighted by previous works. For example, Rachmadi and colleagues(Rachmadi et al., 2018b), for segmenting WMH, extract the brain tissue from the originally acquired MRI, and only input this to the CNN architecture. In addition, perform a three-step intensity normalisation: 1) adjust the maximum grey scale value of the MRI brain to 10 percent of the maximum intensity value, 2) adjust the contrast and brightness of the images such that their histograms are consistent, and 3) normalise the intensities of the resultant images to zero-mean and unit-variance. Guerrero and colleagues, for similar task, used two MRI modalities (Guerrero et al., 2018), which were co-registered, resliced to have 1mmx1mm in-plane voxel size, and normalised their intensities. In general, intensity normalisation, contrast adjustment and removal of background features that could confound the algorithms are necessary for achieving a good segmentation. When multiple MRI sequences or imaging modalities are used, co-registration is also necessary.

#### 1.2.2 Data Augmentation

Training a machine learning model is equivalent to tune its parameters so that it can map a particular input to an output. The number of parameters needed is proportional to the complexity of the task. These parameters can increase if more information is given. The increase in the amount of input data without necessarily meaning an increase in the contextual or semantic data per se is known as data augmentation and has been used in brain image segmentation tasks. Several studies have introduced global spatial information as an additional input to CNN schemes in form of large 2D orthogonal patches downscaled by a factor(de Brebisson and Montana, 2015), integrated with intensity features from image voxels(Van Nguyen et al., 2015), as a number of hand-crafted spatial location features(Ghafoorian et al., 2016), synthetic volume(Steenwijk et al., 2013; Roy et al., 2015), or set of synthetic images that encode spatial information(Rachmadi et al., 2018b) for mentioning some examples. In other words, all input datasets are acquired under a limited set of conditions (e.g. specific MRI scanning protocols, pathology appearance restricted to few examples, etc.). However, our target application may exist in a variety of conditions (e.g. pathologies in different location, scale, brightness, contrasts, shapes). By synthetically generating data to account for these variations without adding irrelevant features, good results might be obtained. A review of the state of the art in medical image analysis concluded that very similar algorithms could achieve different results due to smart data pre-processing and augmentation (Litjens et al., 2017).

#### 1.2.3 Transfer Learning

Transfer learning has become a popular choice for re-purposing machine learning models that have proven useful for particular tasks, by means of either fine-tuning pre-trained models with data of another nature (i.e. domain adaptation transfer learning), or using a pre-trained model as a starting point for a model on a second task of interest (i.e. task adaptation transfer learning). Domain adaptation transfer learning, where data domains in training and testing processes differ, has been applied successfully to brain MRI segmentation tasks. For example, one study improved Support Vector Machines (SVM)’s performance using different distribution of training data(Van Opbroek et al., 2015). Another study pre-trained CNN using natural images for segmentation of neonatal to adult brain images(Xu et al., 2017), and other study pre-trained a CNN for brain brain lesion segmentation using MRI data acquired with other protocols(Ghafoorian et al., 2017). Task adaptation transfer learning has been applied to WMH segmentation, by teaching a CNN to “learn” to detect texture irregularities instead of binary expert-delineated WMH segmentations (Rachmadi et al., 2018a).

### 1.3 Contributions

Our main contributions are to propose and evaluate data augmentation and transfer learning methods for improving the output of a widely used brain lesion segmentation CNN approach, namely DeepMedic, to identify and delineate the ischaemic stroke lesion from MR perfusion imaging.

## 2 METHODS

### 2.1 Data

The ISLES challenge was conceived as a common benchmark for researchers to compare their segmentation algorithms (Maier et al., 2017) for ischaemic stroke lesions. Initially, the first iteration of ISLES (in 2015), included two sub-challenges, namely **S**troke **P**erfusion **ES**timation (SPES) and SISS. The first sub-challenge was about segmenting stroke lesions in the acute phase, whereas the second focused on sub-acute lesions (Maier et al., 2017).

The stroke cases were carefully crafted and included a wide range of lesion variability. Images were obtained in clinical routine, with different amounts of image artifacts and different views (Maier et al., 2017). Also, some subjects suffered from other pathologies that could be mistaken for ischemic stroke lesions. All files are given in uncompressed Neuroimaging Informatics Technology Initiative (NIfTI) format: (*.nii).

ISLES 2017 contains 43 and 32 training and testing acute subjects, respectively. Included MRI sequences are Apparent Diffusion Coefficient (ADC), 4D Perfusion Weighted Image (4DPWI), Mean Transient Time (MTT), relative Cerebral Blood Flow (rCBF), relative Cerebral Blood Volume (rCBV), Time to maximum (Tmax) and Time to peak (TTP). Images from all modalities were skull-stripped, anonymised and individually co-registered.

The Ground Truth (GT) files, which delimit the actual lesion region, were only provided for training subjects, so as to avoid having participants performing fine-tuning on the test data. They were segmented on T2-weighted and Fluid Attenuation Inversion Recovery (FLAIR) sequences after the stroke had stabilised, but these imaging modalities were not provided.

After careful examination, the stroke subjects in the training data were classified into three different stroke subtypes. These are lacunar/subcortical (10 subjects), small cortical (7 subjects) and big cortical/main artery (26 subjects).

### 2.2 Baseline configuration

The baseline CNN model, including its architecture and hyper-parameters, is based on DeepMedic v0.6.1 (Kamnitsas et al., 2017). The architecture used slightly differs from the initial architecture (Kamnitsas et al., 2017). It is illustrated in figure 1, including the addition of residual connections.

**Figure 1.**
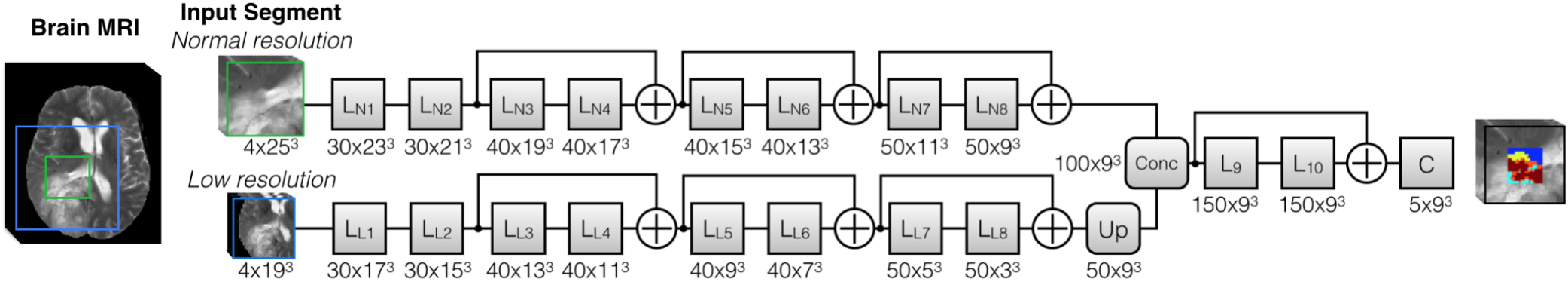
The DeepMedic architecture used, including residual connections.

**Source:** github.com/Kamnitsask/deepmedic

The number of convolutional layers was 8, and the number of feature maps for each were [30, 30, 40, 40, 40, 50, 50]. The kernel size was (3, 3, 3) for all layers. Residual connections in both pathways were also included so that the input of layers [3, 4, 6] was added to the output of layers [4, 6, 8].

The final blocks of the scheme were composed of Fully Connected (FC) layers and a CRF. The number of FC layers was set to two, with 150 feature maps each. The size of the kernels of the first FC layer, which combined the outputs of different scales, was again (3, 3, 3). Additionally, there was a residual connection between the second and first layers, meaning that the input of the first FC layer was added to the output of the second and final FC layer.

The second pathway had an additional parameter that determined the downsampling factor applied to the images fed to the second pathway. Additionally, batch normalization(Ioffe and Szegedy, 2015) was added at the end of each convolutional layer.

The dimension of the training and validation segments were [25, 25, 25] and [17, 17, 17], respectively. The latter was equal to the receptive field of the network. The size of the segments was limited by the available RAM and GPU memory.

The batch size for training, validation and inference were set to 24, 48 and 24, respectively. Dropout(Srivastava et al., 2014) was added in the second FC layer and the final classification layer, both with a rate of 0.5. Weight initialization followed a modified Xavier initialization (Glorot and Bengio, 2010) that accounts for nonlinearities (He et al., 2015). This allows the training of deeper networks and works well with Parametric Rectified Linear Units (PReLU) (He et al., 2015), which were the predefined activation units.

Also, intracranial volume masks were provided to limit the region where samples were extracted from, which in turn saved time and memory. This means that foreground samples were extracted from the GT label mask and background samples extracted from the region inside the subject mask minus the intersection with the label mask. By default, samples were extracted centered in a foreground or background voxel with equal probability.

During training, epochs were divided into subepochs. The number of epochs and subepochs was set to 35 and 20, respectively. For each subepoch, 1000 segments were extracted from up to 50 cases.

The learning rate was decreased exponentially and the momentum linearly increased. The values that had to be reached at the last epoch were 10^−4^ for the former and 0.9 for the latter. The learning rate, initially set to 10^−3^, started to lower at epoch 1. Updating learning rates through training is a way of making sure that convergence is reached and in a reasonable time (Jacobs, 1988; Zeiler, 2012). The learning optimizer was RmsProp(Tieleman and Hinton, 2012), with *ρ* = 0.9 (decay rate) and *ϵ* = 10^−4^ (smoothing term that avoids divisions by zero). RmsProp was combined with Nesterov momentum(Nesterov, 1983), as proposed by (Sutskever et al., 2013). The momentum value was set to *m* = 0.6 and normalized. Additionally, weight decay was also implemented, in the form of L1 and L2 normalization with values *L*1 = 10^−6^ and *L*2 = 10^−4^, respectively.

Also, two “online” (done during training) data augmentation techniques were set by default. The first simply involved reflecting images with a 50% probability with respect to the X axis (from left to right). The second consisted in altering the mean and standard deviation of the images, following the next equation:

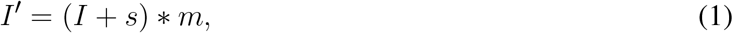

where s (shift) and m (multi) are drawn from Gaussian distributions of (*μ* = 0, *σ* = 0.05) and (*μ* = 1, *σ* = 0.01), respectively.

Finally, due to memory limitations, only three out of the six available channels were used to train the model, namely ADC, MTT and rCBF. In some experiments, rCBF was replaced by rCBV. Only two segmentation classes were considered, foreground, representing the lesion, and background, representing everything else.

### 2.3 Experiments

To evaluate the use of enhancing learning techniques for identifying ischaemic stroke lesions in perfusion imaging data, six experiments were run (i.e. E0-E5) by varying one aspect of the model at a time, such as the type of data or other parameters. This was done in the form of a pipeline, performing pair-wise comparisons. At each stage of the pipeline, two models, with and without a particular change, were compared. The best performing model of each pair-wise comparison proceeded to the next stage, until the best performing model of all experiments was found.

To assess the performance of an experiment, k-fold cross-validation was employed, where *k* = 5. Crossvalidation is essential to give a good estimate of the real performance of an experiment. If cross-validation hadn’t been used, results would have highly depended on the composition of easy/hard cases in each set. For example, if the test set had only been made of easy cases, the performance achieved would have been greater that if they had been difficult cases. Overall, this not only increases the robustness of the results but also the confidence of the decisions related to the changes that have worked best.

#### 2.3.1 Data Pre-processing

Performing adequate pre-processing of the data is essential to maximize the performance of the model. Some of the necessary pre-processing steps were already done by the ISLES organizers, such as coregistering all images per subject setting them to have the same dimension, also per subject, and removing extracranial tissues.

Additional pre-processing involved resampling all images to isotropic (i.e. 1×1×1mm) voxels size, generating intracranial volume masks and normalizing the data to have zero mean and unit variance. The latter is strongly suggested by DeeMedic’s creator as it would substantially affect performance. The intracranial volume masks were generating binarising the TTP images, and applying binary dilation before the resampling to improve the boundaries. Due to memory constraints, all images had to be downsampled with a factor of 0.7 so they could fit in memory.

#### Algorithm 1 Data Pre-processing

~~~
Initialize *dF* = 0.7
**for each** subject **do
    for each** channel **do**
        *resampled_channels* ← *resample*(*channel*)
    **end for**
    *mask* ← *compute_mask*(*channels*)
    *mask* ← *resample*(*mask*)
    *save_image*(*mask*)
    **for each** *resampled_channel* **do**
        *img* ← *normalize*(*resampled_channel, mask*)
        *save_image*(*img*)
    **end for
end for**
~~~

#### 2.3.2 E0 - Baseline Configuration

This experiment (i.e. E0) consisted in training the DeepMedic configuration described previously, with the default parameters using the pre-processed data. It established the baseline results. All future experiments were compared against this or a better performing one. The imaging modalities used as input channels were ADC, MTT and rCBF.

#### 2.3.3 E1 - Data augmentation

We applied the data augmentation method known as intensity variance. It consists in randomly altering the intensity values within the Region of Interest (ROI) or GT region following a Gaussian distribution of mean and variance equal to the ones computed from the intensity values within the region.

The rationale behind this idea was to try to deal with one of the many complications of detecting the ischemic stroke lesion in these types of images: their intensity inhomogeneity. As mentioned by (Maier et al., 2017), the intensity values within the lesion territory can vary significantly. By using a mean and variance based on the already available data, the intensities, while being different from the original, should not be too different so as the lesion is no longer recognizable.

This augmentation was done offline, which means that the altered subjects were created and saved to be fed to the network during training. It was decided to do it this way so as to avoid modifying DeepMedic’s core code, which would in turn become very time consuming. Each new subject is a “clone” of the original, except for the intensity values within the ROI or GT label. All channels had their intensity modified. Algorithm 2 shows how this was done.

##### Algorithm 2 Data augmentation

~~~
Initialize *clones_number* = 1
**for each** subject **do**
    Load *label*
    **for each** *clones_number* **do**
        Initialize *clone_path*
        **for each** *channel* **do**
            *roi* ← *channel*[*nonzero*(*label*)]
            *channel*[*nonzero*(*label*)] ← *gaussian*(*mean*(*roi*), *std*(*roi*))
            *save_image*(*channel, clone_path*)
        **end for
    end for
end for**
~~~

This experiment used the same baseline configuration parameters as E0, with the exception that the data had been augmented. The original 43 subjects had been “cloned”, following the procedure described above. Thus, the total number of available training subjects became 86. However, since validation or testing in augmented subjects is meaningless, only the subjects inside the training set contained clones. Naturally, clones of the validation and test subjects were not part of the training set.

#### 2.3.4 E2 - Transfer learning with error maps

The goal of this experiment was to improve the performance of a pre-trained model (i.e. the best performing model so far), by fine-tuning the model with its error maps (i.e. weighted maps), using them to draw more image segments from difficult regions (i.e. those where errors were bigger).

Fine-tuning is a type of transfer-learning aimed at improving the performance of a network pre-trained for a different-although similar-task to the one the model was originally trained for (Pan et al., 2010). For example, two different tasks can have the same goal and only vary on the information that is provided to complete them. Usually, this technique involves re-training a network while “freezing” the first layers, meaning that their parameters (weights) are kept fixed during training. Each consecutive layer of a CNN generates more complex features from the ones detected in the previous layer. Consequently, the first layers contain simpler features that are common for similar problems, and thus can be “transferred” to a similar task. Then, new data is used to retrain the final layers, tuning the network to improve performance on the new task.

In other words, the aim of fine-tuning is to adapt the network to the small details that make the new task different, which means the learning rate has to account for that by being considerably small compared to the original rate the model was pre-trained with. For that reason, while the learning rate of the initial model was initialized to 10^−3^, the rate for this experiment was 5*x*10^−4^. There are three possible benefits of using transfer learning: a higher start, a higher slope and a higher asymptote(Aytar and Zisserman, 2011). When performing transfer learning, it’s possible that one, two, all or none of these benefits appear.

To improve learning, an adaptive sampling method has been proposed (Berger et al., 2017) for DeepMedic. It consists in extracting more image patches in the regions where the prediction error is bigger, according to error maps generated throughout training. DeepMedic already offers the possibility of using weighted maps for the sampling process, which essentially serves the same function but in a static way (i.e. maps must be generated beforehand and are not updated during training). By using these maps, image segments are extracted more often from those regions where the weights are bigger. Error maps, one per subject and class, were obtained by computing the square error between each voxel of the GT label and the predicted probability map. The probability maps were obtained from the segmented test cases of each fold, meaning that the error maps for all subjects could be computed. These maps were normalized to zero mean and unit variance for homogeneity between subjects.

The paths of the computed error maps were included in different files, one for each class. These files were specified in the configuration parameters, each line representing a subject, which had to be coherent between files. Weighted maps can be defined both for training and validation. Since the goal was to improve the network performance, only error maps for the training cases were provided. In these cases, fine-tuning was performed by retraining the best model so far while extracting more image segments in those regions where errors where bigger, with the aid of pre-computed error maps. All convolutional layers were left frozen, thus only tuning the FC layers.

#### 2.3.5 E3, E4 and E5 - Transfer learning with rCBV

Perfusion parametric maps rCBF and rCBV display different appearance depending on the area under consideration. In the core of the stroke both sequences have substantially low values. However, in the penumbra (i.e. affected but savageable region), while rCBF is slightly reduced, rCBV can be normal or even have higher values compared to normal tissue. Both sequences have been used to segment the stroke (Chen and Ni, 2012).

In this experiment, the best performing model so far is retrained using the ADC, MTT and rCBV as input channels. Recall that until now, models have used the ADC, MTT and rCBF as input channels for training, as defined in the baseline configuration.

The goal of E3 is to make predictions more robust by tuning the weights of the FC layers, similar to experiment E2 in previous section. This would make the network more sensitive to small changes between rCBF and rCBV, which can be crucial to accurately segmenting the stroke.

E4 and E5 are essentially the same as E3 with the exception of the number of frozen layers. E4 has only the first four convolutional layers frozen, whereas E5 has no frozen layers at all. This is useful to also examine the effect of freezing different numbers of layers for the lesion segmentation task.

### 2.4 Post-processing

In order to test whether the predictions of DeepMedic could be further improved, different post-processing techniques were implemented, based on threshold tuning the DeepMedic’s probability output and performing binary morphological operations in the binarised result.

However, before applying any of these techniques, DeepMedic outputs (i.e. predicted lesion and class probability maps) had to be resampled to their corresponding subjects’ original image space so that results could be interpreted in the same dimensional space as the original data. Hence, we resampled all outputs per subject using the inverse affine transformation applied to transform the original images in the ISLES 2017 dataset.

#### 2.4.1 Threshold Tuning

After computing the Receiver Operating Characteristic (ROC) and Precision-Recall (PR) curves it is possible to obtain the optimal threshold to be applied to the DeepMedic probabilistic output, which maximizes the desired metrics. To this end, two threshold tuning procedures, one for each curve, were implemented. It is worth noting that both methods were independent and their results were not combined. Also, both curves were computed using the *Scikit-learn* library.

The first threshold tuning procedure, Threshold Tuning 0 (THT0), consisted in obtaining the point where (*precision * recall*) was maximum. This is the furthest point from the bottom-left corner and thus returns the maximum value for the DSC metric. To compute it, we concatenated the original GT and the probability map of the foreground class of all subjects (separately) to compute the curve, and, then, selected the optimal threshold.

The second procedure, Threshold Tuning 1 (THT1), based on the ROC curve, consisted in obtaining the point where (*TruePositiveRate*(*TPR*) – *FalsePositiveRate*(*FPR*)) was maximum. This represents the furthest point from the bottom-right corner and thus the optimal threshold, giving the maximum value for the Bookmaker Informedness (BM) metric. Again, all subjects’ labels and probability maps were concatenated to compute the curve, and, then, select this threshold.

The goal of both procedures was to obtain the best average threshold for the results from the validation set to apply it to the test set. This was done for all folds independently. This guarantees that the tuning is not performed on the test (i.e. validation) cases, which accounts for a real scenario where the GT for the test cases are not available.

#### 2.4.2 Binary Morphological Operations

Binary morphological operations are mathematical operations used to modify shapes in binary images through a structuring element: a shape to probe the image. Closing is a binary morphological operation that can fill holes in big predicted lesions or join reasonably close small ones to make predictions more robust. It combines two other simpler morphological operations: dilation, which expands shapes in an image, and erosion, which shrinks them. In both cases, the center of the structuring element is placed at every pixel of the image and a decision is made. In the case of dilation, a pixel is set to 1 if there are any pixels equal to one within the shape of the structuring element, otherwise it’s set to zero. Erosion performs the exact opposite operation, a pixel is set to 0 as long as there is any pixel of value 0 within the area covered by the structuring element.

Furthermore, there are two decisions to make regarding this operation: the shape and size of the structuring element and the number of iterations. While the first determines the final output and thus the goodness of the prediction, the second defines the number of times that the closing operation is repeated.

After few experiments, the optimal structuring element was a 3D ball with a radius of 3 voxels, whereas the number of iterations was tuned by selecting the average of the ones that achieved the maximum DSC score on validation cases. This post-processing step was named Filling Holes (FH).

### 2.5 Evaluation

At each state of the post-processing pipeline, multiple performance metrics were computed to compare the predicted segmented lesions with the GT. These metrics were TPR, True Negative Rate (TNR), Positive Predictive Value (PPV), Accuracy (ACC), DSC, Matthews Correlation Coefficient (MCC), and Hausdorff Distance (HD). Being True Positives (TP) the voxels predicted to be positives and identified positives by the configuration evaluated, True Negatives (TN) the voxels predicted to be negatives and identified negatives, False Positives (FP) the voxels predicted to be negatives but identified positives and False negatives (FN),the voxels predicted to be positives but identified negatives, these metrics are defined as follows:

- **TPR:** Also known as *sensitivity* or *recall*, measures the rate of true positives with respect to the number of real positive cases.

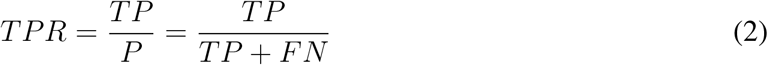
- **TNR:** Also known as *specificity*, measures the rate of true negatives with respect to the number of real negative cases.

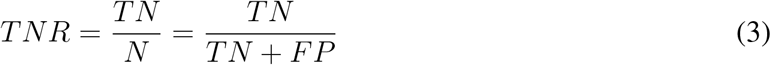
- **PPV:** Also known as *precision*, measures the proportion of true positives with respect to all predicted positives.

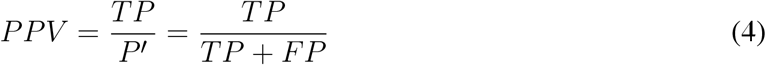
- **ACC:** Is a measure of statistical bias. Represents how close the predictions are from the true values.

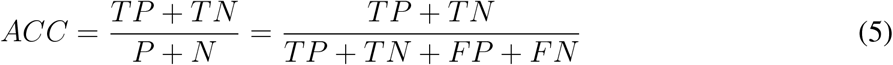
- **DSC:** The Dice similarity coefficient measures the harmonic mean of PPV and TPR. (Landis and Koch, 1977) define the intervals and the associated “strength of agreement”: [< 0.00] (Poor), [0.00 – 0.20] (Slight), [0.21 – 0.40] (Fair), [0.41 – 0.60] (Moderate), [0.61 – 0.80] (Substantial), [0.81 – 1.00] (Almost perfect).

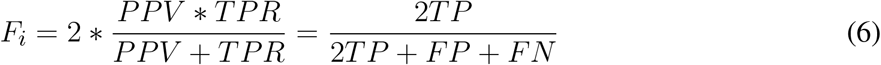
- **MCC:** Also known as the *phi coefficient* or Matthews correlation coefficient, is considered a balanced metric of the quality of binary classification, thus robust to class imbalance. Values range from −1 (perfect negative correlation) to 1 (perfect positive correlation), being 0 equal to random prediction. This metric is considered to be the most meaningful, specially for imbalanced data(Chicco, 2017).

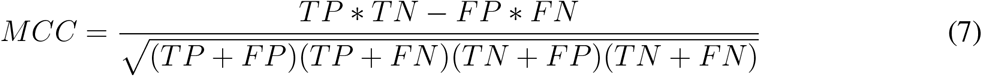
- **HD:** Measures the distance between two subsets. *A_S_* and *B_S_* are equivalent to P (real true cases) and P′ (predicted true cases), and *d*(·) is the euclidean distance between two points.

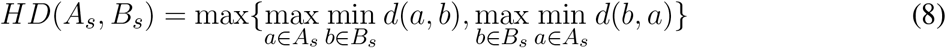

Since k-fold cross-validation was employed, these metrics were averaged per fold and also between folds. This means that performance metrics were available per subject (both for the validation and test sets’ subjects of every fold), per fold and per experiment. Performance curves, known as precision PPV vs. recall TPR, error bar and Bland-Altman(Bland et al., 1986) plots were also produced. In addition, the DeepMedic plotting script was slightly modified to generate the progress of metrics such as accuracy or DSC on training and validation sets through the different epochs.

## 3 RESULTS

### 3.1 Segmentation Performance during Training

The segmentation performance for validation and training sets during the training process is shown in figure 2. The DSC coefficient was stable after improving during few epochs. On the other hand, sensitivity (i.e. TPR) improved at first but then worsened and remained stable. Mean accuracy and specificity, while being very high, did not account for the imbalanced nature of the data.

**Figure 2.**
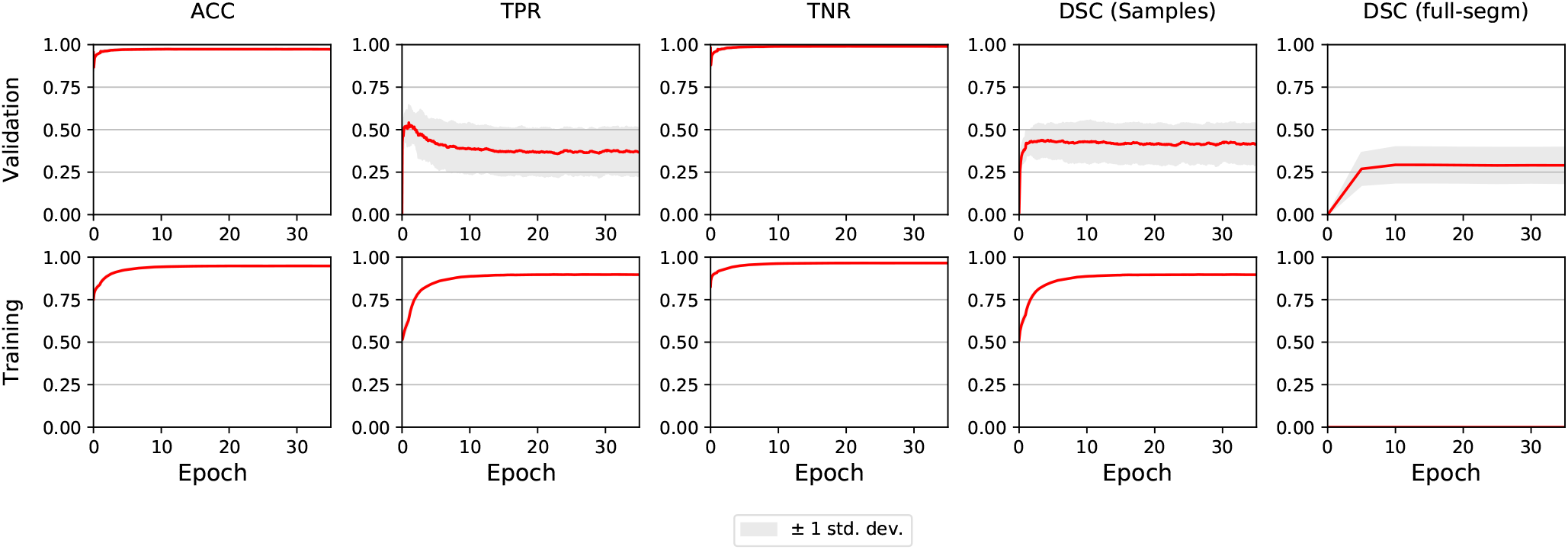
E0 - Segmentation metrics of validation and train subjects during training. The graphs shown are the averages of all 5 folds. The light grey area illustrates ±1 standard deviation. Full segmentation on training cases was not performed by DeepMedic, reason why the lower-right graph is empty.

In E1, sensitivity took more time to reach its peak compared to E0, but when it stabilised the asymptote was slightly higher. Also, while DSC behaved similarly to E0, it also achieved higher values. In E2-E5, the metrics for the first epoch had the same value as for the last epoch in E1, and did not improve throughout the training process.

### 3.2 Baseline Segmentation Performance

Figure 3, shows the error bars for each metric, post-processing step and lesion category for E0. TPR was highly variable for small stroke lesions, regardless of whether they were lacunar or cortical, especially after the THT0 and FH post-processing steps. THT1 produced consistently worse results in terms of accuracy for small stroke lesions, despite achieving higher TPR (i.e. sensitivity). The segmentation of big cortical/main artery stroke lesions was considerably better than those for the other stroke subtypes.

**Figure 3.**
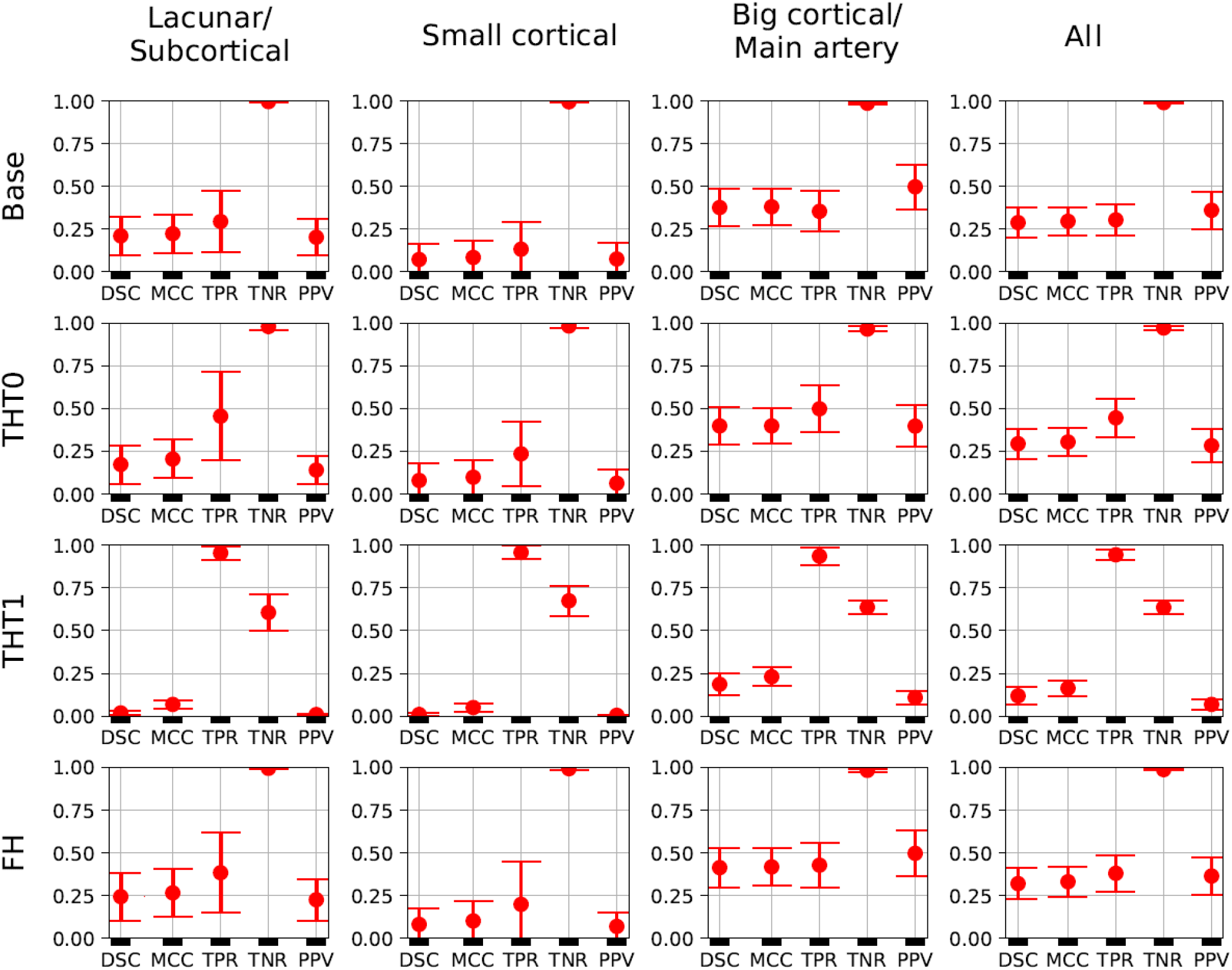
E0 - Error bars. Each metric for each post-processing step and lesion category is presented. A fourth column, representing all subjects, is also included. Each marker represent the mean value, and the upper and lower limits represent the 95% confidence interval. The metrics shown are: Dice similarity coefficient (DSC), Matthews correlation coefficient (MCC), True positive rate (TPR), True negative rate (TNR), and Positive predicted value (PPV).

The Bland-Altman plot showing the volumetric agreement between the GT and the results from E0 after each post-processing step can be seen in figure 4. THT1 produced the worst results in terms of volumetric agreement regardless of the stroke subtype, considerably inflating the stroke lesion volume. This method for selecting the optimal threshold for binarising the probabilistic stroke lesion maps obtained, overestimated the stroke lesion size in general.

**Figure 4.**
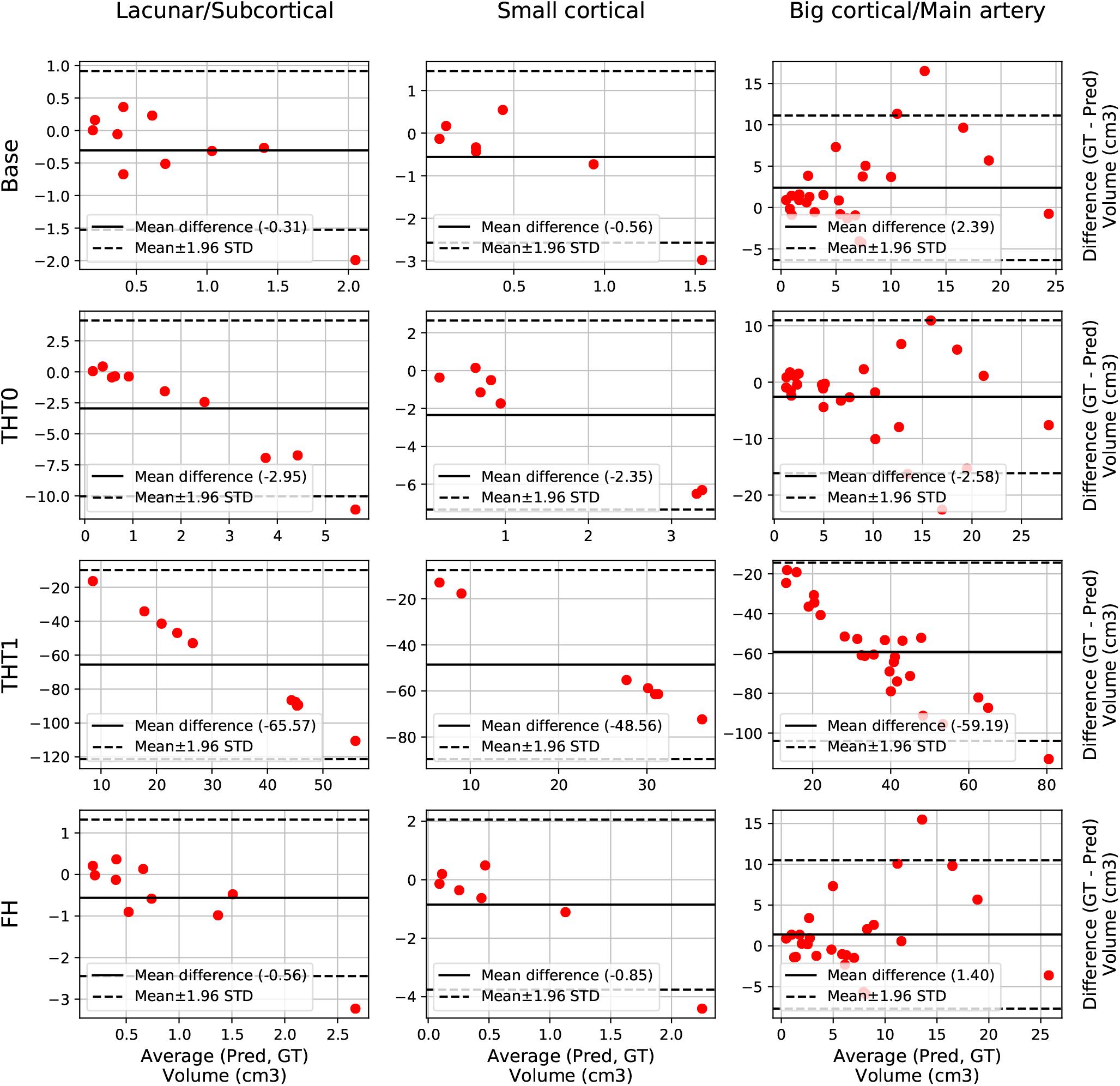
E0 - Volume Bland-Altman analysis. Each lesion category (lacunar/subcortical, small cortical and big cortical) and post-processing step (THT0, THT1, FH and base) are included. Each point represents one subject. The black line is the mean difference, whereas the black dotted-line represents the limits of agreement, computed as mean±1.96 Standard deviation (STD). The x axis is the average volume between the predicted segmentation and the ground truth, whereas the y label is the difference.

### 3.3 Experiments’ Results

E1 was the best performing model, with an average DSC of 0.34 after applying FH. This proves the efficacy of using the data augmentation method selected (i.e. intensity variance). It also proves the importance of performing post-processing tasks, such as THT0 and FH, instead of simply focusing on pre-processing and then relying on the output of the network.

Table 1 and figure 5 contain a summary of all experiments. E1 was superior to E0 and the rest experiments yielded results close to E1, but they were not able to improve it. E4 and E5 are not shown because their results were very similar to E3 but slightly inferior. In general, the transfer learning approaches (E2-E5) evaluated did not improve the accuracy in the results.

**Table 1.** Summary of the main metrics for all experiments (i.e. E0-E3). Average metrics from the base prediction and all post-processing steps are shown. These are: Threshold tuning 0 (THT0), Threshold tuning 1 (THT1) and Filling holes (FH). The metrics shown are: Dice similarity coefficient (DSC), Hausdorff distance (HD), Matthews correlation coefficient (MCC), True positive rate (TPR), True negative rate (TNR), and Positive predicted value (PPV).

|  | Post-proc | DSC | HD | MCC | TPR | TNR | PPV |
| --- | --- | --- | --- | --- | --- | --- | --- |
| **E0** | **Base** | 0.29 | 62.22 | 0.30 | 0.30 | 0.99 | 0.36 |
|  | **THT0** | 0.29 | 72.83 | 0.30 | 0.45 | 0.97 | 0.28 |
|  | **THT1** | 0.12 | 99.62 | 0.16 | 0.94 | 0.64 | 0.07 |
|  | **FH** | 0.32 | 59.47 | 0.33 | 0.38 | 0.99 | 0.36 |
| **E1** | **Base** | 0.32 | 49.89 | 0.32 | 0.34 | 0.99 | 0.38 |
|  | **THT0** | 0.31 | 72.33 | 0.33 | 0.49 | 0.97 | 0.30 |
|  | **THT1** | 0.13 | 100.29 | 0.18 | 0.96 | 0.65 | 0.07 |
|  | **FH** | 0.34 | 47.85 | 0.35 | 0.40 | 0.99 | 0.39 |
| **E2** | **Base** | 0.31 | 48.48 | 0.32 | 0.34 | 0.99 | 0.38 |
|  | **THT0** | 0.31 | 71.42 | 0.33 | 0.48 | 0.97 | 0.30 |
|  | **THT1** | 0.13 | 100.19 | 0.18 | 0.96 | 0.68 | 0.08 |
|  | **FH** | 0.33 | 46.74 | 0.35 | 0.40 | 0.99 | 0.38 |
| **E3** | **Base** | 0.30 | 57.37 | 0.31 | 0.36 | 0.99 | 0.36 |
|  | **THT0** | 0.31 | 66.37 | 0.32 | 0.42 | 0.98 | 0.32 |
|  | **THT1** | 0.12 | 99.94 | 0.17 | 0.97 | 0.63 | 0.07 |
|  | **FH** | 0.33 | 53.94 | 0.34 | 0.42 | 0.99 | 0.36 |

**Figure 5.**
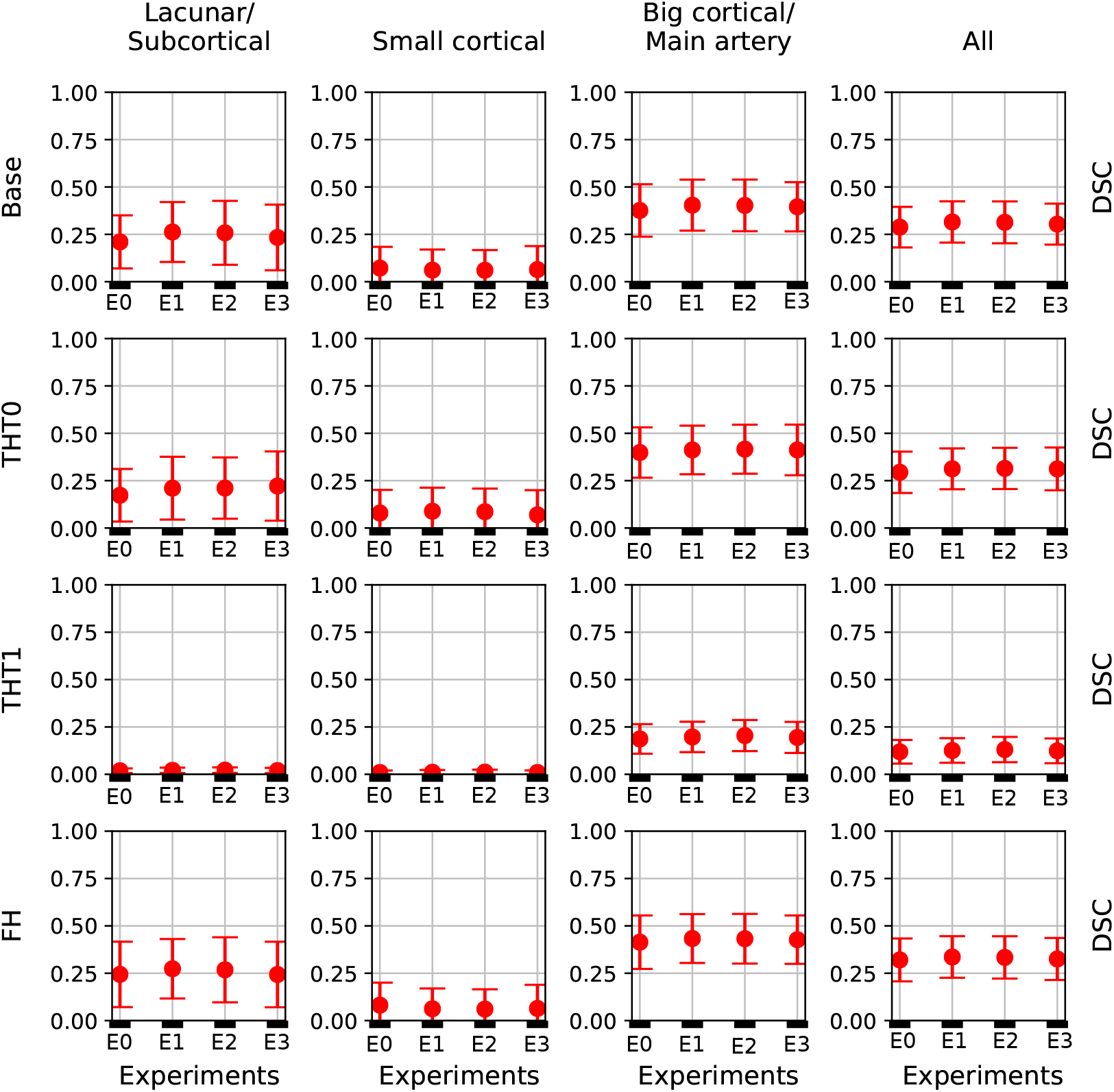
DSC error bars of all experiments for the base prediction and FH and each lesion category.

Table 1 shows the key metrics of each experiment both for all post-processing steps. On average, FH performed best. PPV and consequently DSC were the metrics that determined the best performing model.

Figure 5 depicts the DSC error bars for all post-processing steps and lesion categories. Big cortical lesions were easier to segment than the rest (i.e. small lesions).

Additionally, figure 6 shows the precision-recall curves for all experiments. Results are very different depending on the cases that fall in each fold. This is a clear sign of the heterogeneous nature of the data and the inability of the network to generalising well. Also from these graphs, results from E1 are slightly superior to E0 and similar to E2. Interestingly, while E3 produced the worst results, its predictions were the least heterogeneous (i.e. the curves are more closer to each other than in any other experiment).

**Figure 6.**
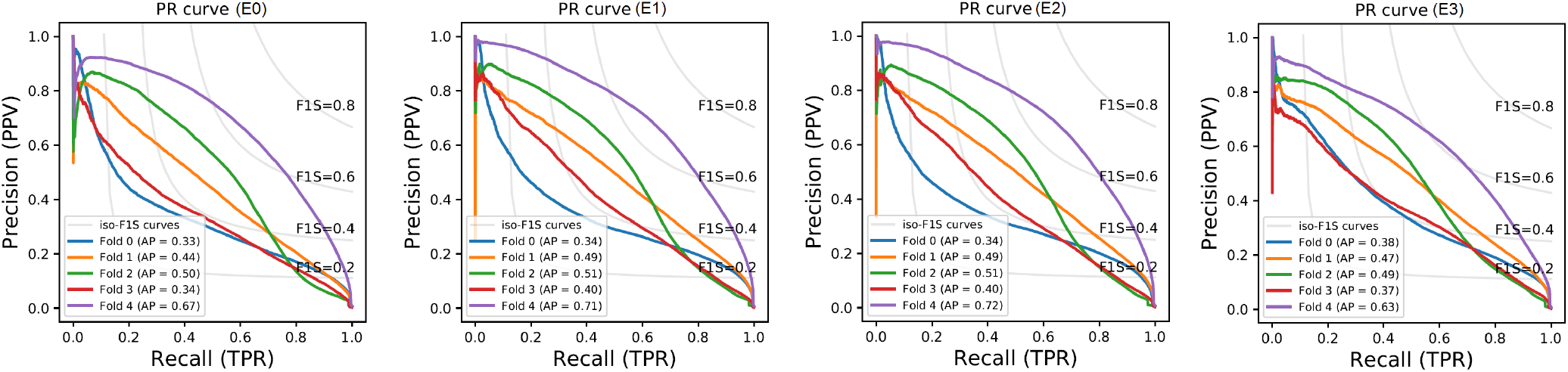
Performance curves of E0-E3. The grey lines indicate the iso-F1 Score (F1S) curves, the value of DSC for each point in the graph. The Average Precision (AP) metrics are also included.

The winner (Choi et al., 2017) of the ISLES 2017 challenge, achieved 0.31 DSC and 103.64 HD when the final results were published in September of 2017, but since then the challenge has remained open. Consequently, more participants have joined the challenge and the current top performer, as of the time of writing this manuscript, achieved 0.36 DSC and 29.37 HD.

To perform a fair comparison between our E1 and the current state of the art performance, E1 was retrained using all train data for training and tested on the unlabeled test set of the challenge. FH was then applied to the predicted lesions using the average number of iterations in E1 and the results uploaded to the SMIR web page^2^.

E1 achieved 0.29 DSC and 49.75 HD on the test set, as reported by the SMIR web page. This value is inferior to the 0.34 DSC achieved in the E1 experiment and also to the current first position of the challenge. This difference could be because of the fact that either the network or the number of iterations for FH computed in E1 were not able to generalize well on the test data.

### 3.4 Visual Evaluation of the Results

Figures 7, 8 and 9 show the results from E1 for representative axial slices superimposed in the ADC image, from three subjects randomly selected from each category. In general, stroke lesion predictions were better in E1, but not by a large margin, and these figures, overall, exemplify the results obtained.

**Figure 7.**
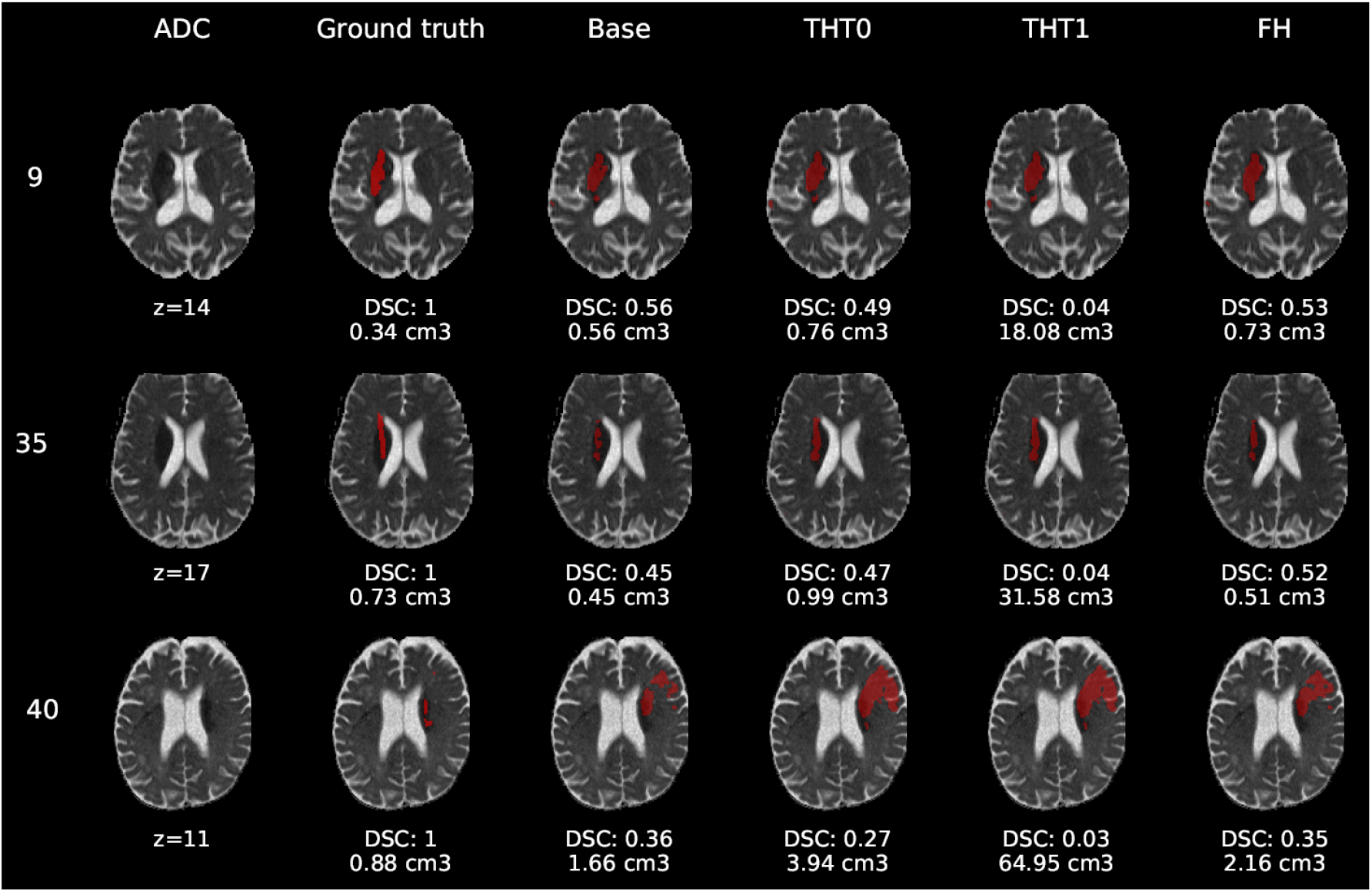
E1 - Visual segmentation comparison of lacunar/subcortical lesions. The examples include the predicted lesions after each post-processing step. Images are 2D slices, their cut coordinate in the z axis is included, as well as the volume of each segmentation and the DSC achieved.

**Figure 8.**
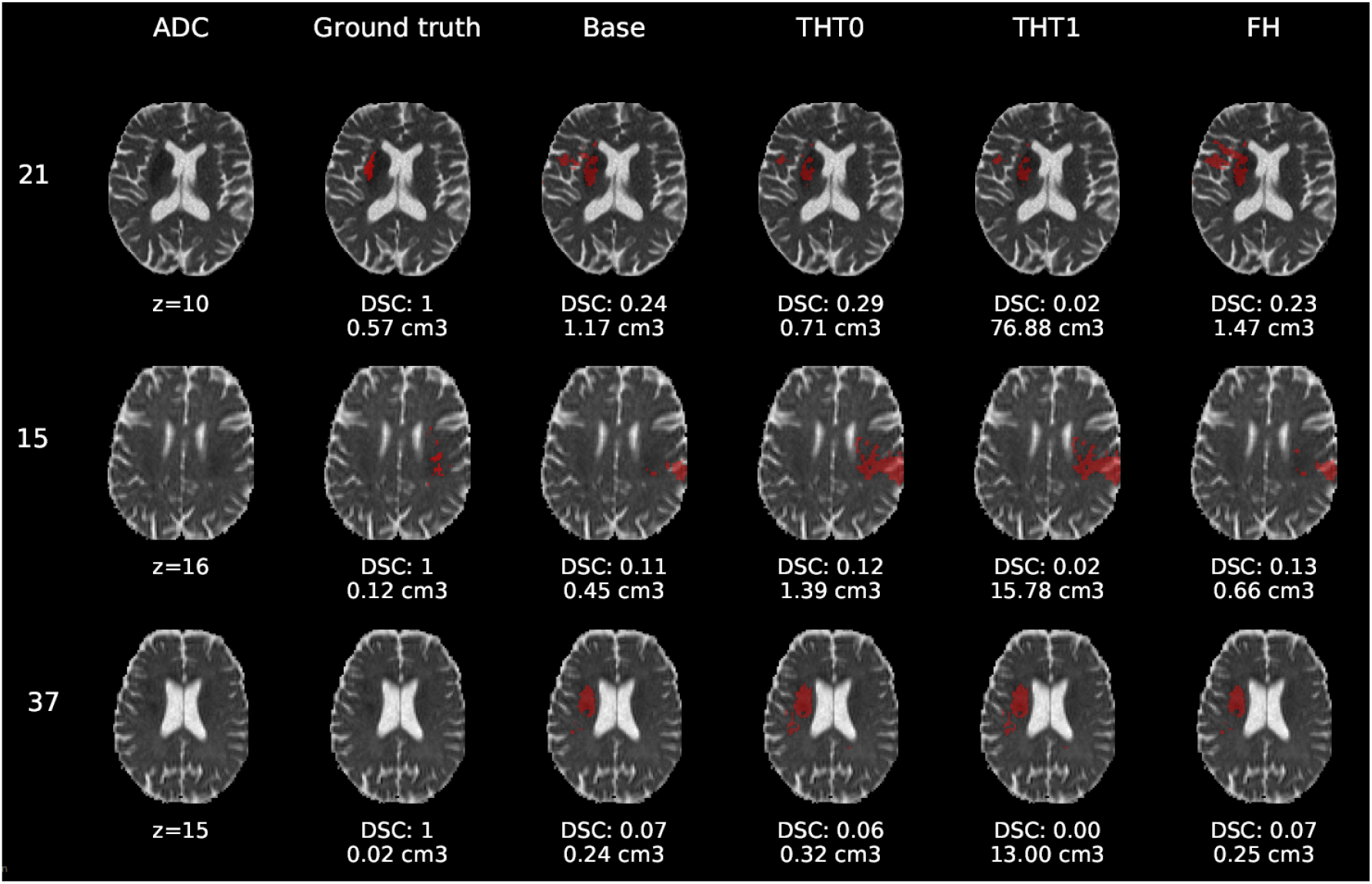
E1 - Visual segmentation comparison of small cortical lesions. The examples include the predicted lesions after each post-processing step. Images are 2D slices, their cut coordinate in the z axis is included, as well as the volume of each segmentation and the DSC achieved.

**Figure 9.**
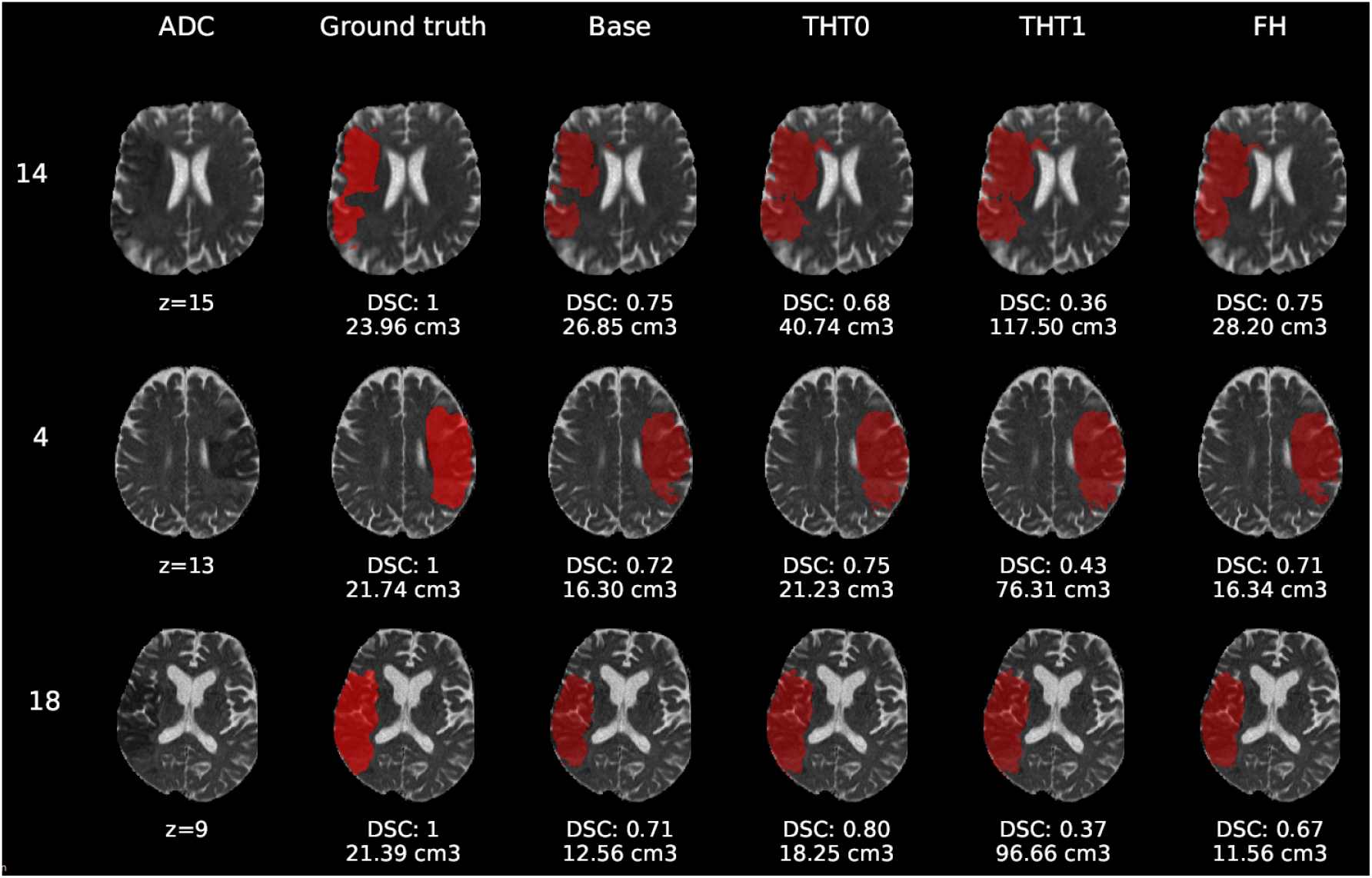
E1 - Visual segmentation comparison of big cortical lesions. The examples include the predicted lesions after each post-processing step. Images are 2D slices, their cut coordinate in the z axis is included, as well as the volume of each segmentation and the DSC achieved.

Compared to E0, some cases were better segmented, but this was not always the case. For example, the stroke lesion prediction for subject 9 (lacunar infarct) achieved a DSC score of 0.45 in E0, whereas in E1 it achieved 0.56. However, for subject 21 (small cortical infarct), the DSC score for E0 was 0.26, whereas in E1 it was 0.24, i.e. a slightly worse score. In general, E1’s DSC was 10.34% better than E0’s and 6.25% for FH. Most results were visually very similar. Also, in E1, post-processing steps (i.e. THT0, THT1, FH) did not improve results as much as they did in E0.

The GT, obtained from the structural T2-weighted images, not always includes the whole regions with restricted diffusion (i.e. dark regions in the ADC map). Contrastingly, in cases of large strokes, it includes the cerebrospinal fluid in the sulci. For cases in which the GT extent agrees with the region of restricted diffusion, the results are better (e.g. cases 9 and 32).

Visually, results obtained applying THT1 to the DeepMedic’s output does not appear to be disparately wrong compared to those obtained applying THT0 and/or FH.

## 4 DISCUSSION

The model that used data augmentation had the best performance, achieving an average DSC score of 0.34 for the test cases after applying FH. This was a reasonable outcome considering that the network clearly suffered from overfitting, for which data augmentation is a well-known remedy.

Also, of all post-processing steps evaluated, FH produced the best improvements on average over the base prediction by the network. The second best was THT0, which in some cases surpassed FH. The results from applying THT1, although worst in terms of accuracy, were not visually very different.

Despite the enhancing learning strategy proposed slightly improved the segmentation results in the majority of cases, our results are still suboptimal. We used the default configuration, batch size, learning rate and activation functions of a CNN scheme designed to segment tumours from structural MRI sequences. Also, instead of pre-training the network with data of similar nature, but a varied, larger dataset, and fined-tune it with this ISLES 2017 dataset, we directly trained it with a subset from the latter. Therefore, overfitting was still a problem even with data augmentation. Reducing it could be achieved by modifying the number of layers and the size of kernels, and thus the number of network parameters. It could also be remedied by using data from other challenges, or even past iterations of ISLES that also contain the same sequences for segmenting the stroke lesion. Moreover, the learning rate schedule should lower the learning rate at predefined epochs. We used the DeepMedic’s default without prior training the model to determine when it would be more convenient to lower the learning rate, and the schedule was set to exponential decrease. Further work should try to lower the learning rate only when necessary.

Despite the limitations previously mentioned, the GT used should be put into question. As the examples selected show, it did not accurately cover the region of restricted diffusion in the ADC images, underestimating it mainly for small infarcts and overestimating in cases of large infarcts, including regions of cerebrospinal fluid in the sulci. The GT was generated using the structural T2-weighted images (i.e. including FLAIR), not provided. The mismatch between structural, diffusion and perfusion MRI modalities is well-known (Motta et al., 2015; Chen and Ni, 2012; Straka et al., 2010).

Precisely, the perfusion/diffusion mismatch has been reported to provide a practical and approximate measure of the tissue at risk, being used to identify acute stroke patients that could benefit from reperfusion therapies. Clinical studies also show that early abnormality on diffusion-weighted imaging can overestimate the infarct core by including part of the tissue “at risk”, and the abnormality on perfusion weighted imaging overestimates this “at risk” tissue by including regions of benign tissue with reduced blood perfusion (Chen and Ni, 2012).

The diffusion/fluid attenuated inversion recovery (DWI/FLAIR) mismatch is also well known. Together with the perfusion/diffusion mismatch it is recognised as an MRI marker of evolving brain ischemia. A clinical trial that examined whether the DWI/FLAIR mismatch was independently associated with the diffusion/perfusion mismatch or not, concluded that in the presence of the latter, the DWI/FLAIR pattern could indicate a shorter time between the scan and the last time the tissue seen was normal (Wouters et al., 2015). The CNN scheme evaluated does not take into account the time from the stroke onset - information not provided.

Finally, the types of infarcts were not evenly represented in the dataset. The large cortical strokes were predominant, which could explain the bias in the results favouring the cases when the stroke was of this subtype. The involvement of personnel with relevant clinical knowledge in the generation of datasets to be used for developing algorithms aimed to clinical research would be advisable in the future.

## AUTHOR CONTRIBUTIONS

C.U.P.M., M.C.V.H., M.F.R., and T.K. conceived and presented the idea. C.U.P.M. and M.C.V.H. planned the experiments. C.U.P.M. carried out the experiments. All authors provided critical feedback and analysis, and contributed for the manuscript.

## FUNDING AND ACKNOWLEDGEMENTS

This study was partially funded by the Indonesia Endowment Fund for Education (LPDP) of Ministry of Finance, Republic of Indonesia (M.F.R.), Row Fogo Charitable Trust (Grant No. BRO-D.FID3668413)(M.C.V.H.), the European Union Horizon 2020 [PHC-03-15, project No 666881, ‘SVDs@Target’] (M.C.V.H.), and the UK Biotechnology and Biological Sciences Research Council (BBSRC) through the International Partnership Award BB/P025315/1 (M.C.V.H.).

## DATA AVAILABILITY STATEMENT

The dataset used in this study can be found in the ISLES Challenge 2015 repository (www.isles-challenge.org/ISLES2015/). The code that corresponds with the experiments described and analysed in this manuscript can be found in https://github.com/CarlosUziel/ischleseg.

## Footnotes

1 www.isles-challenge.org/ISLES2015/

2 www.smir.ch

## REFERENCES

Aytar, Y. and Zisserman, A. (2011). Tabula rasa: Model transfer for object category detection. In Computer Vision (ICCV), 2011 IEEE International Conference on (IEEE), 2252–2259

Berger, L., Hyde, E., Cardoso, J., and Ourselin, S. (2017). An adaptive sampling scheme to efficiently train fully convolutional networks for semantic segmentation. arXiv preprint arXiv:1709.02764

Bland, J. M., Altman, D. G., et al. (1986). Statistical methods for assessing agreement between two methods of clinical measurement. lancet 1, 307–310

Brosch, T., Tang, L. Y., Yoo, Y., Li, D. K., Traboulsee, A., and Tam, R. (2016). Deep 3d convolutional encoder networks with shortcuts for multiscale feature integration applied to multiple sclerosis lesion segmentation. IEEE transactions on medical imaging 35, 1229–1239

Chen, F. and Ni, Y.-C. (2012). Magnetic resonance diffusion-perfusion mismatch in acute ischemic stroke: An update. World journal of radiology 4, 63

Chen, L., Papandreou, G., Schroff, F., and Adam, H. (2017). Rethinking atrous convolution for semantic image segmentation. CoRR abs/1706.05587

Chen, L.-C., Papandreou, G., Kokkinos, I., Murphy, K., and Yuille, A. L. (2018a). Deeplab: Semantic image segmentation with deep convolutional nets, atrous convolution, and fully connected crfs. IEEE transactions on pattern analysis and machine intelligence 40, 834–848

Chen, L.-C., Zhu, Y., Papandreou, G., Schroff, F., and Adam, H. (2018b). Encoder-decoder with atrous separable convolution for semantic image segmentation. arXiv preprint arXiv:1802.02611

Chicco, D. (2017). Ten quick tips for machine learning in computational biology. BioData mining 10, 35

Choi, Y., Kwon, Y., Paik, M. C., and Joon, B. (2017). Ischemic stroke lesion segmentation with convolutional neural networks for small data. ISLES 2017 Challenge

de Brebisson, A. and Montana, G. (2015). Deep neural networks for anatomical brain segmentation. In Proceedings of the IEEE Conference on Computer Vision and Pattern Recognition Workshops. 20–28

Fantini, S., Sassaroli, A., Tgavalekos, K. T., and Kornbluth, J. (2016). Cerebral blood flow and autoregulation: current measurement techniques and prospects for noninvasive optical methods. Neurophotonics 3, 031411

Ghafoorian, M., Karssemeijer, N., van Uden, I. W., de Leeuw, F.-E., Heskes, T., Marchiori, E., et al. (2016). Automated detection of white matter hyperintensities of all sizes in cerebral small vessel disease. Medical physics 43, 6246–6258

Ghafoorian, M., Mehrtash, A., Kapur, T., Karssemeijer, N., Marchiori, E., Pesteie, M., et al. (2017). Transfer learning for domain adaptation in mri: Application in brain lesion segmentation. In International Conference on Medical Image Computing and Computer-Assisted Intervention (Springer), 516–524

Glorot, X. and Bengio, Y. (2010). Understanding the difficulty of training deep feedforward neural networks. In Proceedings of the thirteenth international conference on artificial intelligence and statistics. 249–256

Guerrero, R., Qin, C., Oktay, O., Bowles, C., Chen, L., Joules, R., et al. (2018). White matter hyperintensity and stroke lesion segmentation and differentiation using convolutional neural networks. NeuroImage: Clinical 17, 918–934

He, K., Zhang, X., Ren, S., and Sun, J. (2014). Spatial pyramid pooling in deep convolutional networks for visual recognition. In european conference on computer vision (Springer), 346–361

He, K., Zhang, X., Ren, S., and Sun, J. (2015). Delving deep into rectifiers: Surpassing human-level performance on imagenet classification. In Proceedings of the IEEE international conference on computer vision. 1026–1034

He, K., Zhang, X., Ren, S., and Sun, J. (2016). Deep residual learning for image recognition. In Proceedings of the IEEE conference on computer vision and pattern recognition. 770–778

Ioffe, S. and Szegedy, C. (2015). Batch normalization: Accelerating deep network training by reducing internal covariate shift. arXiv preprint arXiv:1502.03167

Jacobs, R. A. (1988). Increased rates of convergence through learning rate adaptation. Neural networks 1, 295–307

Kamnitsas, K., Ledig, C., Newcombe, V. F., Simpson, J. P., Kane, A. D., Menon, D. K., et al. (2017). Efficient multi-scale 3d cnn with fully connected crf for accurate brain lesion segmentation. Medical image analysis 36, 61–78

Krizhevsky, A., Sutskever, I., and Hinton, G. E. (2012). Imagenet classification with deep convolutional neural networks. In Advances in neural information processing systems. 1097–1105

Landis, J. R. and Koch, G. G. (1977). The measurement of observer agreement for categorical data. biometrics, 159–174

Litjens, G., Kooi, T., Bejnordi, B. E., Setio, A. A. A., Ciompi, F., Ghafoorian, M., et al. (2017). A survey on deep learning in medical image analysis. Medical image analysis 42, 60–88

Lucas, C. and Heinrich, M. P. (2017). 2d multi-scale res-net for stroke segmentation. ISLES 2017 Challenge

Maier, O., Menze, B. H., von der Gablentz, J., Häni, L., Heinrich, M. P., Liebrand, M., et al. (2017). Isles 2015-a public evaluation benchmark for ischemic stroke lesion segmentation from multispectral mri. Medical image analysis 35, 250–269

Milletari, F., Navab, N., and Ahmadi, S.-A. (2016). V-net: Fully convolutional neural networks for volumetric medical image segmentation. In 3D Vision (3DV), 2016 Fourth International Conference on (IEEE), 565–571

Motta, M., Ramadan, A., Hillis, A. E., Gottesman, R. F., and Leigh, R. (2015). Diffusion–perfusion mismatch: an opportunity for improvement in cortical function. Frontiers in neurology 5, 280

Nesterov, Y. E. (1983). A method for solving the convex programming problem with convergence rate o (1/k^^^ 2). In Dokl. Akad. Nauk SSSR. vol. 269, 543–547

Pan, S. J., Yang, Q., et al. (2010). A survey on transfer learning. IEEE Transactions on knowledge and data engineering 22, 1345–1359

Petrella, J. R. and Provenzale, J. M. (2000). Mr perfusion imaging of the brain: techniques and applications. American Journal of roentgenology 175, 207–219

Rachmadi, M. F., del C. Valdés-Hernández, M., and Komura, T. (2018a). Transfer learning for task adaptation of brain lesion assessment and prediction of brain abnormalities progression/regression using irregularity age map in brain mri. In PRedictive Intelligence in MEdicine, eds. I. Rekik, G. Unal, E. Adeli, and S. H. Park (Cham: Springer International Publishing), 85–93

Rachmadi, M. F., del C. Valdés-Hernández, M., Agan, M. L. F., Perri, C. D., and Komura, T. (2018b). Segmentation of white matter hyperintensities using convolutional neural networks with global spatial information in routine clinical brain mri with none or mild vascular pathology. Computerized Medical Imaging and Graphics 66, 28 – 43. doi:https://doi.org/10.1016/j.compmedimag.2018.02.002

Roth, H. R., Lu, L., Seff, A., Cherry, K. M., Hoffman, J., Wang, S., et al. (2014). A new 2.5 d representation for lymph node detection using random sets of deep convolutional neural network observations. In International Conference on Medical Image Computing and Computer-Assisted Intervention (Springer), 520–527

Roy, P. K., Bhuiyan, A., Janke, A., Desmond, P. M., Wong, T. Y., Abhayaratna, W. P., et al. (2015). Automatic white matter lesion segmentation using contrast enhanced flair intensity and markov random field. Computerized Medical Imaging and Graphics 45, 102–111

Sermanet, P., Eigen, D., Zhang, X., Mathieu, M., Fergus, R., and LeCun, Y. (2013). Overfeat: Integrated recognition, localization and detection using convolutional networks. arXiv preprint arXiv:1312.6229

Srivastava, N., Hinton, G., Krizhevsky, A., Sutskever, I., and Salakhutdinov, R. (2014). Dropout: a simple way to prevent neural networks from overfitting. The Journal of Machine Learning Research 15, 1929–1958

Steenwijk, M. D., Pouwels, P. J., Daams, M., van Dalen, J. W., Caan, M. W., Richard, E., et al. (2013). Accurate white matter lesion segmentation by k nearest neighbor classification with tissue type priors (knn-ttps). NeuroImage: Clinical 3, 462–469

Straka, M., Albers, G. W., and Bammer, R. (2010). Real-time diffusion-perfusion mismatch analysis in acute stroke. Journal of Magnetic Resonance Imaging 32, 1024–1037

Sudre, C. H., Li, W., Vercauteren, T., Ourselin, S., and Cardoso, M. J. (2017). Generalised dice overlap as a deep learning loss function for highly unbalanced segmentations. In Deep Learning in Medical Image Analysis and Multimodal Learning for Clinical Decision Support (Springer). 240–248

Sutskever, I., Martens, J., Dahl, G., and Hinton, G. (2013). On the importance of initialization and momentum in deep learning. In International conference on machine learning. 1139–1147

Tieleman, T. and Hinton, G. (2012). Lecture 6.5-rmsprop: Divide the gradient by a running average of its recent magnitude. COURSERA: Neural networks for machine learning 4, 26–31

Van Nguyen, H., Zhou, K., and Vemulapalli, R. (2015). Cross-domain synthesis of medical images using efficient location-sensitive deep network. In International Conference on Medical Image Computing and Computer-Assisted Intervention (Springer), 677–684

Van Opbroek, A., Ikram, M. A., Vernooij, M. W., and De Bruijne, M. (2015). Transfer learning improves supervised image segmentation across imaging protocols. IEEE transactions on medical imaging 34, 1018–1030

Wouters, A., Dupont, P., Ringelstein, E. B., Norrving, B., Chamorro, A., Grond, M., et al. (2015). Association between the perfusion/diffusion and diffusion/flair mismatch: data from the axis2 trial. Journal of Cerebral Blood Flow & Metabolism 35, 1681–1686

Xu, Y., Géraud, T., and Bloch, I. (2017). From neonatal to adult brain mr image segmentation in a few seconds using 3d-like fully convolutional network and transfer learning. In Image Processing (ICIP), 2017 IEEE International Conference on (IEEE), 4417–4421

Zeiler, M. D. (2012). Adadelta: an adaptive learning rate method. arXiv preprint arXiv:1212.5701

